# Endothelial SMAD1/5 signaling couples angiogenesis to osteogenesis during long bone growth

**DOI:** 10.1101/2023.01.07.522994

**Authors:** Annemarie Lang, Andreas Benn, Angelique Wolter, Tim Balcaen, Joseph Collins, Greet Kerckhofs, An Zwijsen, Joel D. Boerckel

**Affiliations:** Departments of Orthopaedic Surgery and Bioengineering, University of Pennsylvania, Philadelphia, PA, United States; Charité – Universitätsmedizin Berlin, corporate member of Freie Universität Berlin, Humboldt-Universität zu Berlin, and Berlin Institute of Health, Department of Rheumatology and Clinical Immunology, Berlin, Germany; Center for Molecular and Vascular Biology, Department of Cardiovascular Sciences, KU Leuven, Belgium; VIB-KU Leuven Center for Brain & Disease Research, KU Leuven, 3000 Leuven, Belgium; Institute of Animal Welfare, Animal Behavior and Laboratory Animal Science, Department of Veterinary Medicine, Freie Universität Berlin, Berlin, Germany; Biomechanics lab, Institute of Mechanics, Materials and Civil Engineering, UCLouvain, Louvain-la-Neuve, Belgium; Pole of Morphology, Institute of Experimental and Clinical Research, UCLouvain, Brussels, Belgium; Molecular Design and Synthesis, Department of Chemistry, KU Leuven, Leuven, Belgium; Department of Materials Engineering, KU Leuven, Heverlee, Belgium; Prometheus, Division for Skeletal Tissue Engineering, KU Leuven, Leuven, Belgium

**Author notes:** These authors contributed equally (AL, AB). To whom correspondence should be addressed: An Zwijsen, Department of Cardiovascular Sciences, Campus Gasthuisberg, O&N1, Herestraat 49 box 911, 3000 Leuven, Belgium, Joel D. Boerckel, McKay Orthopaedic Research Laboratory, 376A Stemmler Hall, 36th Street & Hamilton Walk, Philadelphia, PA 19104, USA.

## Abstract

Skeletal development depends on coordinated angiogenesis and osteogenesis. Bone morphogenetic proteins direct bone development by activating SMAD1/5 signaling in osteoblasts. However, the role of SMAD1/5 in skeletal endothelium is unknown. Here, we found that endothelial cell-conditional SMAD1/5 depletion in juvenile mice caused metaphyseal and diaphyseal hypervascularity, resulting in altered cancellous and cortical bone formation. SMAD1/5 depletion induced excessive sprouting, disrupting the columnar structure of the metaphyseal vessels and impaired anastomotic loop morphogenesis at the chondro-osseous junction. Endothelial SMAD1/5 depletion impaired growth plate resorption and, upon long term depletion, abrogated osteoprogenitor recruitment to the primary spongiosa. Finally, in the diaphysis, endothelial SMAD1/5 activity was necessary to maintain the sinusoidal phenotype, with SMAD1/5 depletion inducing formation of large vascular loops, featuring elevated endomucin expression, ectopic tip cell formation, and hyperpermeability. Together, endothelial SMAD1/5 activity sustains skeletal vascular morphogenesis and function and coordinates growth plate remodeling and osteoprogenitor recruitment dynamics during bone growth.

## Introduction

The development of the skeleton depends on spatiotemporally coordinated blood vessel morphogenesis and bone formation. In development, multicellular patterning is coordinated by morphogens (*1*). During bone development, morphogens of the bone morphogenetic protein (BMP) family are principal regulators of osteogenesis and signal via intracellular effector proteins, including SMAD1 and SMAD5 (SMAD1/5) (*2, 3*). Despite the abundance of BMP ligands and the coordinated coupling of angiogenesis and osteogenesis during bone growth, the role of SMAD1/5 signaling in skeletal endothelium has not been studied (*4, 5*).

Diverse BMP ligands are abundant during bone development and are expressed by a variety of cell types, including skeletal endothelial cells (*7–12*). The functions of these morphogens, and that of their downstream SMAD signaling, has been studied extensively in skeletal-lineage cells, resulting in FDA-approved therapies for bone formation and regeneration (*12–15*). Multiple studies in various established angiogenesis models, including embryonic development, the mouse retina, and the zebrafish (*11, 16–18*), demonstrate that SMAD signaling is also important to endothelial cell function. Previously, we demonstrated that, during embryonic mouse development, SMAD1/5 synergize with Notch signaling to balance the selection of tip and stalk cells in developmental vascular sprouting (*19*). Further, we observed that endothelial cell-specific depletion of SMAD1/5 during early postnatal retinal angiogenesis reduced the number of tip cells, caused hyperdensity of the vascular plexus, and induced arteriovenous malformations (*20*). However, the role of SMAD1/5 in long-bone blood vessels, which develop in this particularly BMP-rich niche, is unknown.

Endothelial cells exhibit remarkable genetic and phenotypic heterogeneity, which enables diverse and specialized vascular functions. The growing skeleton contains two primary types of blood vessels: “type H” and “type L”, marked by their high- and low-expression, respectively, of the endothelial (surface) markers, CD31 and endomucin (EMCN). Type H vessels (CD31^hi^EMCN^hi^) have columnar/looping structure, form by bulging angiogenesis, and functionally couple angiogenesis to osteogenesis at the growth plate during endochondral long bone growth (*4*). Type L vessels (CD31^low^EMCN^low^) have sinusoidal structure, form by sprouting angiogenesis in the bone marrow, and functionally couple with hematopoiesis in the bone marrow. Both type H and type L vessels can be targeted for genetic manipulation using Cdh5-Cre^ERT2^-mediated recombination.

Here, we performed Cdh5-conditional homozygous depletion of both SMAD1 and SMAD5 to evaluate the role of BMP-SMAD signaling in the formation, maintenance, and function of the type H and type L vessels and their regulation of postnatal bone growth. We performed both short-term (7 day) and long-term (14 day) SMAD1/5 depletion from the endothelium of both juvenile and adolescent mice. Juvenile (P21-P35) and adolescent (P42-P56) ages were selected to evaluate vascular morphogenesis during periods of rapid and modest bone growth, respectively. We show that endothelial SMAD1/5 signaling regulates both type H and type L vessel morphogenesis, maintenance, and function, and couples angiogenesis to growth plate remodeling and osteoprogenitor cell maintenance to control skeletal growth. These findings provide insights into how endothelial BMP-SMAD1/5 signaling contributes to bone formation and homeostasis and may contribute to a better understanding of clinical applications of BMPs for vascularized bone regeneration (*21*).

## Results

### SMAD1/5 restricts vessel volume and width during vascular growth in long bones

To study the role of endothelial SMAD1/5 signaling in morphogenesis of the long bone vasculature, we generated inducible, endothelial cell-conditional (Cdh5-Cre^ERT2^) *Smad1/5* double knockout mice (SMAD1/5^iΔEC^), which were compared to Cre-negative littermate controls (SMAD1/5^WT^). Mice were injected daily with tamoxifen at postnatal day 21-23 (P21-P23) and tibia samples were taken at P28 (**Fig. 1A**). Efficiency of Cre-recombination in ECs was shown previously (*20*) and verified by reduction in phospho-SMAD1/5-positive ECs in the bone marrow (**Fig. S1**). We used contrast-enhanced microfocus X-ray computed tomography (CECT) analysis of the tibia to visualize and quantify the metaphyseal and bone marrow vasculature in 3D. Endothelial cell-conditional SMAD1/5 depletion at weaning resulted in significantly dilated vessels with disrupted morphology in both metaphyseal and diaphyseal vessels within one week post-knockout. Specifically, SMAD1/5 depletion significantly increased the relative vessel volume (VV/TV) in both the metaphysis and diaphysis (*metaphysis p = 0.037; diaphysis p < 0.001;* **Fig. 1C, E**) and increased the relative vessel surface in the diaphysis (*p = 0.001*) (**Fig. 1D, F**). Measurement of vascular linear density (V.Li.Dn) indicated that SMAD1/5 depletion did not alter vessel number in the metaphysis (**Fig. 1G**), but significantly elevated vessel number in the diaphysis (*p = 0.022;* **Fig. 1K**). SMAD1/5 depletion significantly elevated the mean vessel width in both the metaphysis and diaphysis by 46.4% and 55.6%, respectively (*p < 0.001*), reducing the frequency of smaller capillaries (<0.04 mm) and increasing the frequency of larger vessels (**Fig. 1H, I, L, M**). SMAD1/5 depletion did not significantly alter vascular separation (i.e., spacing between vessels) in the metaphysis, but reduced vascular separation in the diaphysis (*p < 0.001;* **Fig. 1J, N**). These data demonstrate a critical role of postnatal endothelial SMAD1/5 signaling in shaping and maintaining the 3D morphology of both metaphyseal and diaphyseal vessels.

**Figure 1.**
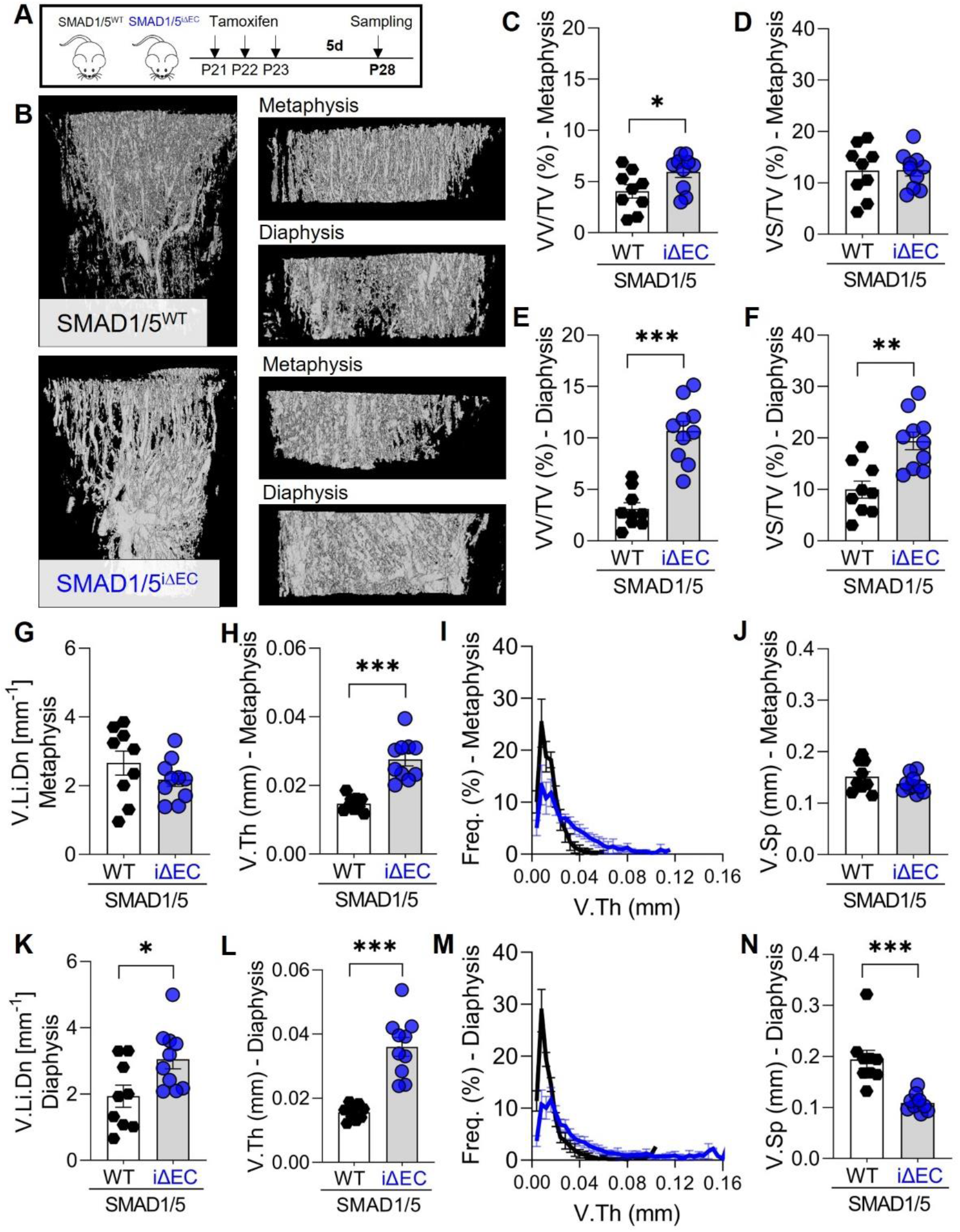
Endothelial SMAD1/5 depletion after weaning increased metaphyseal and diaphyseal vascularity. (**A**) Tamoxifen treatment scheme. Mice were injected postnatal day 19-21 (P19-21) and samples were collected at P28. (**B**) CECT-based 3D rendering visualizing vessels (P28). Quantitative CECT-based structural analysis (P28; n^WT^= 9; n^iΔEC^= 10) of (**C**) relative vessel volume (VV/TV) and (**D**) surface (VS/TV) in the metaphysis or (**E, F**) diaphysis, respectively. (**G**, **K**) vessel linear density (V.Li.Dn), (**H**, **L**) mean vessel thickness (V.Th) and (**I**, **M**) frequency, as well as (**J**, **N**) vascular separation (V.Sp) in metaphysis and diaphysis, respectively. Bar graphs show mean ± SEM and individual data points. Two-sample t-test or Mann Whitney U test (V.Sp, diaphysis) was used to determine the statistical significance; p-values are indicated with *\*p < 0.05; **p < 0.01; ***p < 0.001*.

### Endothelial SMAD1/5 activity directs cortical bone formation during long bone growth

Postnatal long bone growth occurs through both endochondral ossification at the growth plate and cortical bone maturation (*22*). To determine the role of endothelial SMAD1/5 activity in cancellous and cortical bone formation, we examined the bone morphometrical parameters of the metaphyseal and diaphyseal regions of the tibia using μCT (**Fig. 2A**). EC-specific SMAD1/5 depletion significantly reduced trabecular number (Tb.N; *p= 0.047*), but did not alter bone volume fraction (BV/TV), trabecular thickness (Tb.Th), or separation (Tb.Sp) (**Fig. 2B-E**). SMAD1/5 depletion significantly reduced cortical bone area (Ct. Ar) and the cortical area fraction (Ct.Ar/Tt.Ar) in the diaphysis (*p = 0.042* and *p= 0.013*, respectively; **Fig. 2F-H**) and decreased cortical porosity (Ct.Po) (*p= 0.014;* **Fig. 2I**). These findings indicate that endothelial SMAD1/5 signaling directs cortical bone maturation and highlight the importance of vessel morphogenesis in long bone growth.

**Figure 2.**
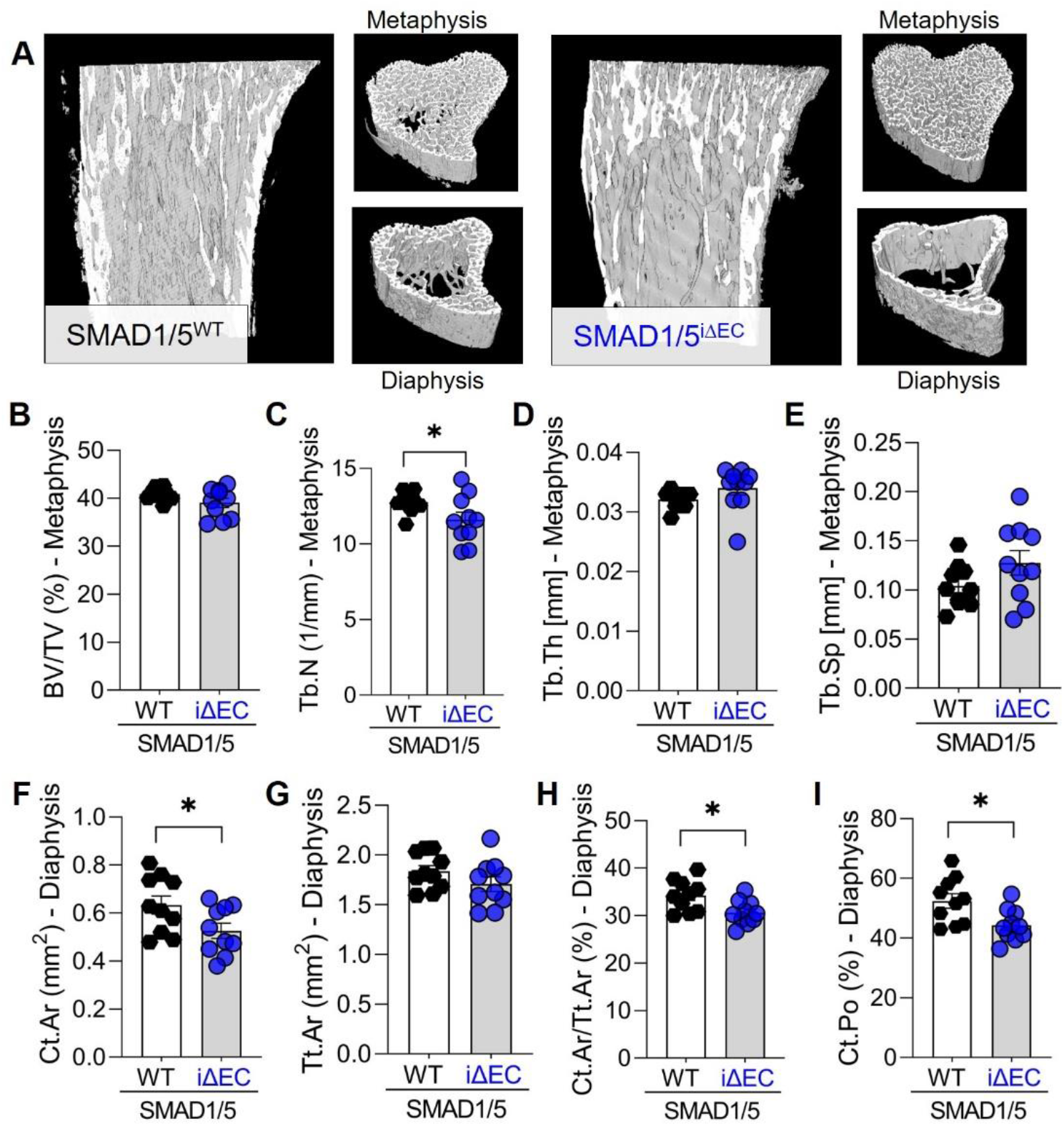
Endothelial SMAD1/5 depletion after weaning decreased diaphyseal formation. (**A**) μCT-based 3D rendering of the proximal tibia at P28. Quantitative μCT-based structural analysis (P28; n^WT^= 9; n^iΔEC^= 10) of (**B**) bone volume fraction (BV/TV), (**C**) trabecular number (Tb.N), (**D**) trabecular thickness (Tb.Th) and (**E**) trabecular separation (Tb.Sp) in the metaphysis or (**F**) cortical bone area (Ct.Ar), (**G**) total cross-sectional area (Tt.Ar), (**H**) cortical area fraction (Ct.Ar/Tt.Ar) and (**I**) cortical porosity (Ct.Po) in the diaphysis. Bar graphs show mean ± SEM and individual data points. Two-sample t-test was used to determine the statistical significance; p-values are indicated with *\*p < 0.05*.

### Angiogenic-osteogenic coupling in the metaphysis requires endothelial SMAD1/5 activity

Type H vessels couple angiogenesis and bone formation during endochondral ossification. These specialized capillaries exhibit columnar structure, terminate at the growth plate in anastomotic arches, and associate with Osterix-expressing (OSX^+^) osteoprogenitor cells (*4*). To determine the role of postnatal SMAD1/5 signaling in type H vessel morphogenesis and angiogenic-osteogenic coupling, we performed histomorphometry for type H vessels (CD31^hi^EMCN^hi^; **Fig. 3A, B**) and OSX^+^ cells (**Fig. 4**) in the metaphysis. EC-specific SMAD1/5 depletion induced aberrant branching of the type H vessels, impairing their columnar structure, and reduced the number of anastomotic arches (mean difference = 3.8 ± 1.2 arches/mm; *p = 0.02*) adjacent to the growth plate (**Fig. 3C**), but did not significantly alter EMCN^+^ and CD31^+^ area in the metaphysis (**Fig. 3D, E**). These data demonstrate a role of endothelial SMAD1/5 signaling in short term-morphogenesis of the type H vessels.

**Figure 3.**
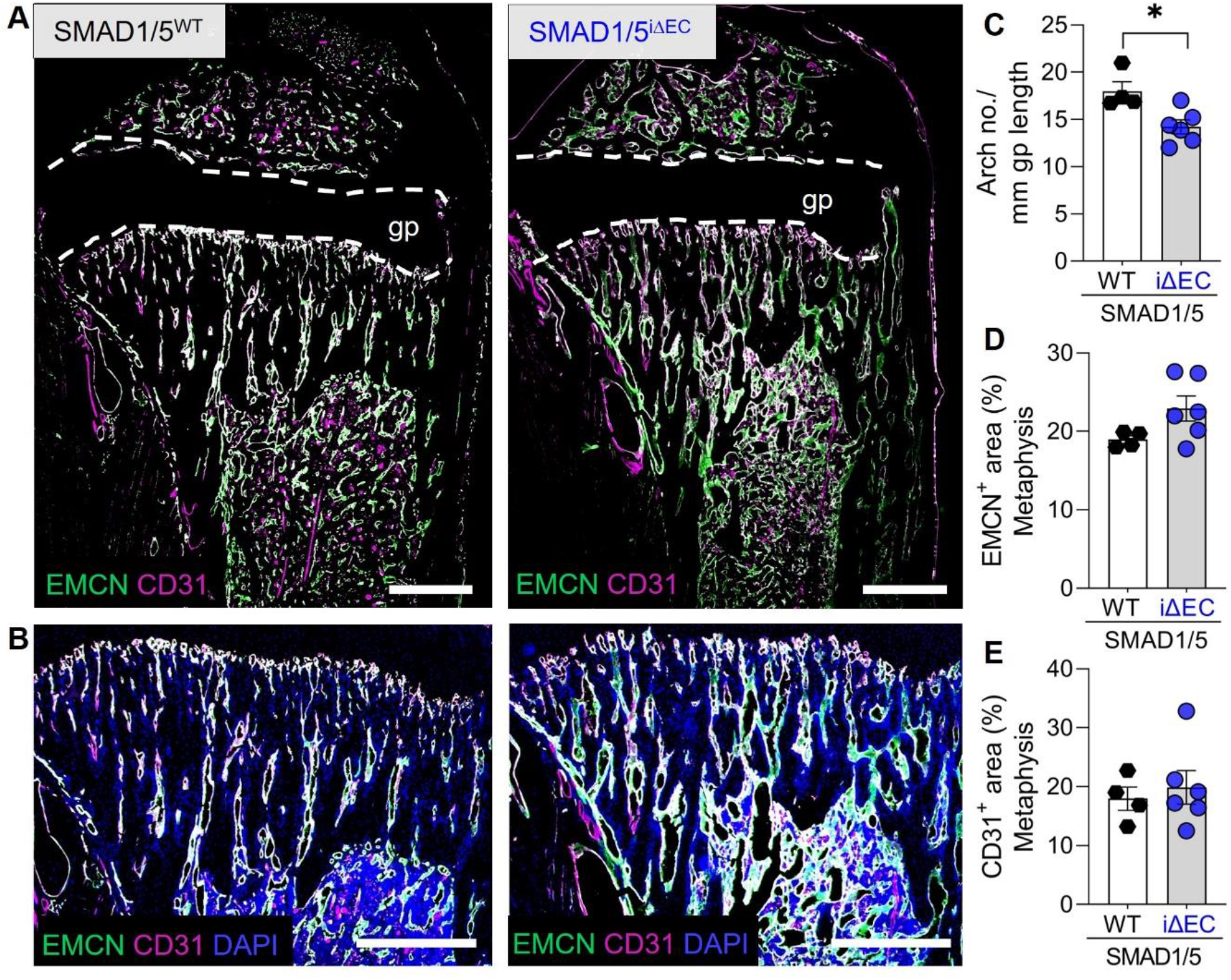
Endothelial SMAD1/5 activity contributes to type H vessel morphology. (**A**) Representative images of EMCN and CD31 staining in the tibial metaphyseal and diaphyseal area (P28; n^WT^= 4; n^iΔEC^= 6). (**B**) Magnifications of (**A**) showing EMCN, CD31 and DAPI staining in the metaphyseal area (P28; n^WT^= 4; n^iΔEC^= 6). Quantification of (**C**) arch number, (**D**) EMCN^+^ and (**E**) CD31^+^ areas (P28; n^WT^= 4; n^iΔEC^= 6). gp – growth plate. Bar graphs show mean ± SEM and individual data points. Two-sample t-test was used to determine the statistical significance; p-values are indicated with \**p < 0.05*. All scale bars indicate 500 μm.

**Figure 4.**
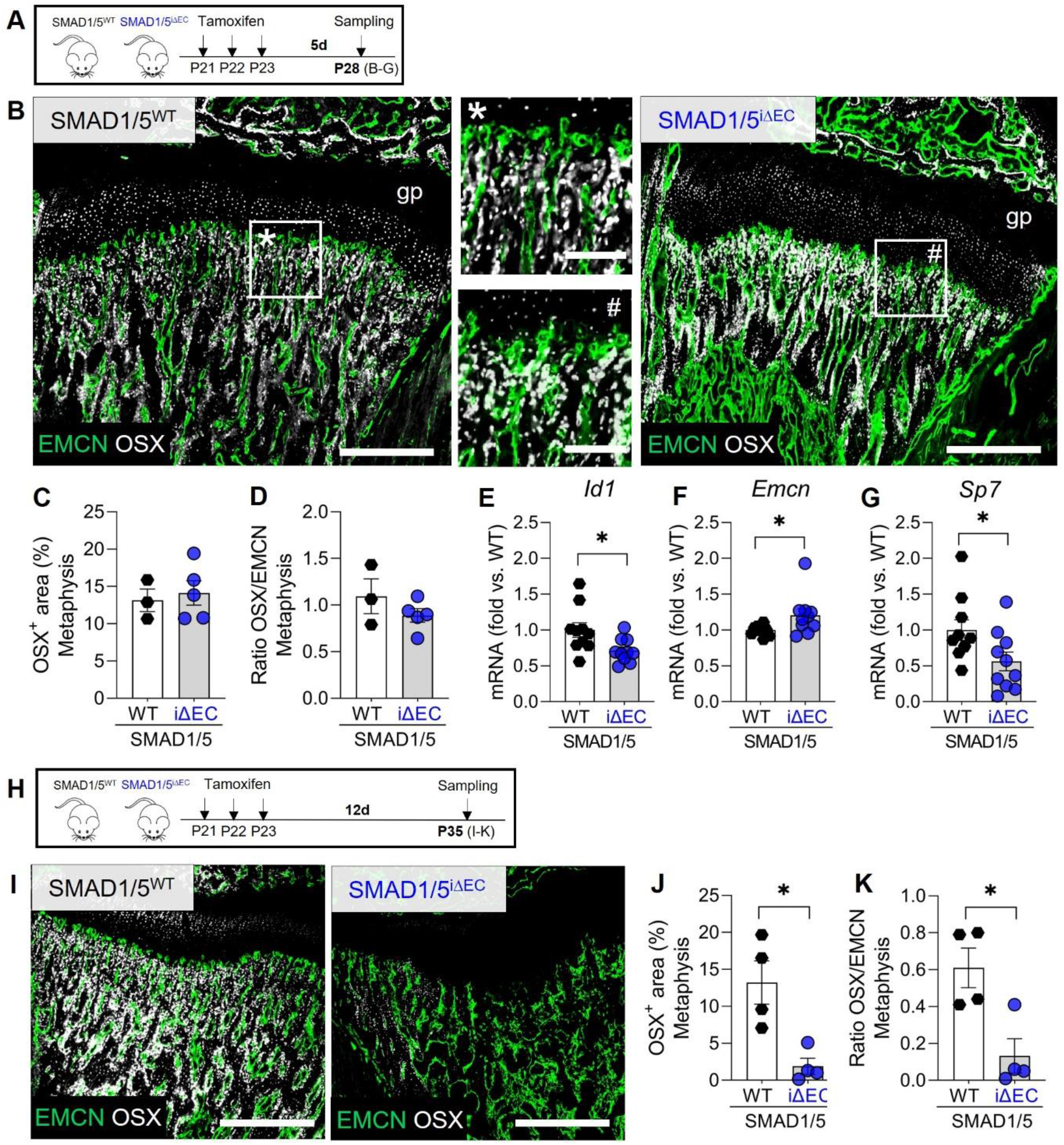
Endothelial SMAD1/5 ensures vascular co-localization of active osteoprogenitors in the metaphyseal area. (**A**) Tamoxifen treatment scheme. Mice were injected postnatal day 19-21 (P19-21) and samples were collected at P28. (**B**) Representative images of EMCN and OSX staining in the metaphyseal area (P28; n^WT^= 4; n^iΔEC^= 6). gp – growth plate.) Quantification of (**C**) OSX^+^ cell area and (**D**) ratio of OSX/EMCN (P28; n^WT^= 4; n^iΔEC^= 6). mRNA expression analysis of (**E**) *Id1*, (**F**) *Emcn* and (**G**) *Sp7* in the epi-/metaphysis (P28; n= 10). (**H**) Tamoxifen treatment scheme. Mice were injected postnatal day 19-21 (P19-21) and samples were collected at P35. (**I**) Representative images of EMCN and OSX staining in the metaphyseal area (P35; n^WT^= 4; n^iΔEC^= 4). Quantification of (**J**) OSX^+^ cell area and (**K**) ratio of OSX/EMCN (P35; n^WT^= 4; n^iΔEC^= 4). Bar graphs show mean ± SEM and individual data points. Two-sample t-test or Mann Whitney U test (*Emcn* RNA expression) was used to determine the statistical significance; p-values are indicated with *\*p < 0.05*. All scale bars indicate 500 μm (**B**, **I**) or 125 μm (magnifications **B**).

Type H vessels physically associate with OSX^+^ osteoprogenitor cells and couple angiogenesis to osteogenesis during postnatal bone growth (*4*). Therefore, we next evaluated the effects of endothelial SMAD1/5 depletion on OSX^+^ osteoprogenitors dynamics in the metaphysis by quantifying OSX^+^ cells at two time points after tamoxifen-induced depletion (P28 vs. P35; 7d vs. 14d post-tamoxifen injection). As above, we first evaluated OSX^+^ cells at P28 (7d after first tamoxifen injection) (**Fig. 4A**). EC-specific depletion of SMAD1/5 did not significantly alter OSX^+^ cells or OSX/EMCN ratio in the metaphysis at 7 days post-depletion (P28; **Fig. 4A-D; Fig. S2**). Bulk gene expression analysis was performed on metaphyseal and epiphyseal tissue to evaluate expression of the canonical SMAD1/5-target gene, *Id1* (*23*). As expected, *Id1* expression was significantly lower in the meta-/epiphysis of SMAD1/5^iΔEC^ mice (*p = 0.02;* **Fig. 4E**). Consistent with immunostaining for EMCN (**Fig. 3D**), EC-specific depletion of SMAD1/5 increased *Emcn* mRNA abundance (*p = 0.03;* **Fig. 4F**). Notably, *Sp7* (OSX) mRNA was significantly reduced (44% lower, *p = 0.04*) by endothelial SMAD1/5 deactivation at P28 (**Fig. 4G**). These data suggested that the angiogenic-osteogenic coupling between type H vessels and surrounding osteoprogenitors was beginning to be altered at this early timepoint (7 days post-depletion; P28). We therefore performed an additional OSX^+^ cell analysis at P35 (14 days after tamoxifen injection; **Fig. 4H, I**). By two-weeks post tamoxifen, EC-specific SMAD1/5 depletion significantly and markedly decreased the abundance of OSX^+^ osteoprogenitors (*p= 0.011*) and OSX/EMCN ratio (*p= 0.015*) in the metaphysis (**Fig. 4J, K**). Together, these data indicate that endothelial SMAD1/5 activity regulates type H vessel morphogenesis and is required for maintenance of osteoprogenitor cells in the metaphysis, functionally coupling angiogenesis and osteogenesis during juvenile bone growth.

### Loss of metaphyseal vessel integrity results in accumulation of hypertrophic chondrocytes in the growth plate

The anastomotic arches of the type H capillaries function not only to support network connectivity and osteoprogenitor mobilization, but also actively degrade the hypertrophic cartilage to enable endochondral ossification (*24*). Therefore, we next asked whether the disruption of the type H vessel structures caused by endothelial SMAD1/5 depletion affected the morphogenesis and remodeling of the hypertrophic cartilage at the chondro-osseous junction. For the experimental design, we chose the same procedure as for the osteoprogenitor analysis to address growth plate remodeling dynamics. Thus, we investigated growth plate changes at P28 (7d after first tamoxifen injection; **Fig. 5A**) and also P35 (14 days after first tamoxifen injection; **Fig. 5E**). Depletion of SMAD1/5 activity in ECs did not significantly alter cell morphology, thickness, or hypertrophic chondrocyte fraction in the growth plate at 7 days post-tamoxifen (**Fig. 5B, Fig. S3**). Consistently, SMAD1/5 depletion did not alter metaphyseal mRNA expression of *Mmp9, Ctsk, Adamts1* and *Timp1* at 7 days post-tamoxifen. However, by P35, 14 days post-tamoxifen, EC-specific SMAD1/5 depletion resulted in dysmorphogenesis of the anastomotic arches at the chondro-osseous junction (**Fig. 5D**; arrows) and a significant enlargement of the hypertrophic zone (hz) of the growth plate (**Fig. 5D, E**). Quantification of the total growth plate size indicated that reduced endothelial SMAD1/5 activity did not induce a general enlargement of the total growth plate (*p= 0.1*) but a shift of zonal distribution with a significant increase in the relative COL10^+^ area (27% increase; *p= 0.0003;* **Fig. 5E**). Chondrocytes occupy lacunae in the extracellular matrix which can be counted in parallel to DAPI^+^ nuclei for assessment of growth plate cellularity. The number of DAPI^+^ cells and chondrocyte lacuna in the COL10^+^ area was slightly increased, suggesting an increase in cellular quantity rather than a volumetric enlargement (**Fig. 5F**). Together, these data establish the necessity of ongoing SMAD1/5 signaling in maintenance of type H anastomotic arch-mediated resorption of hypertrophic cartilage and growth plate remodeling.

**Figure 5.**
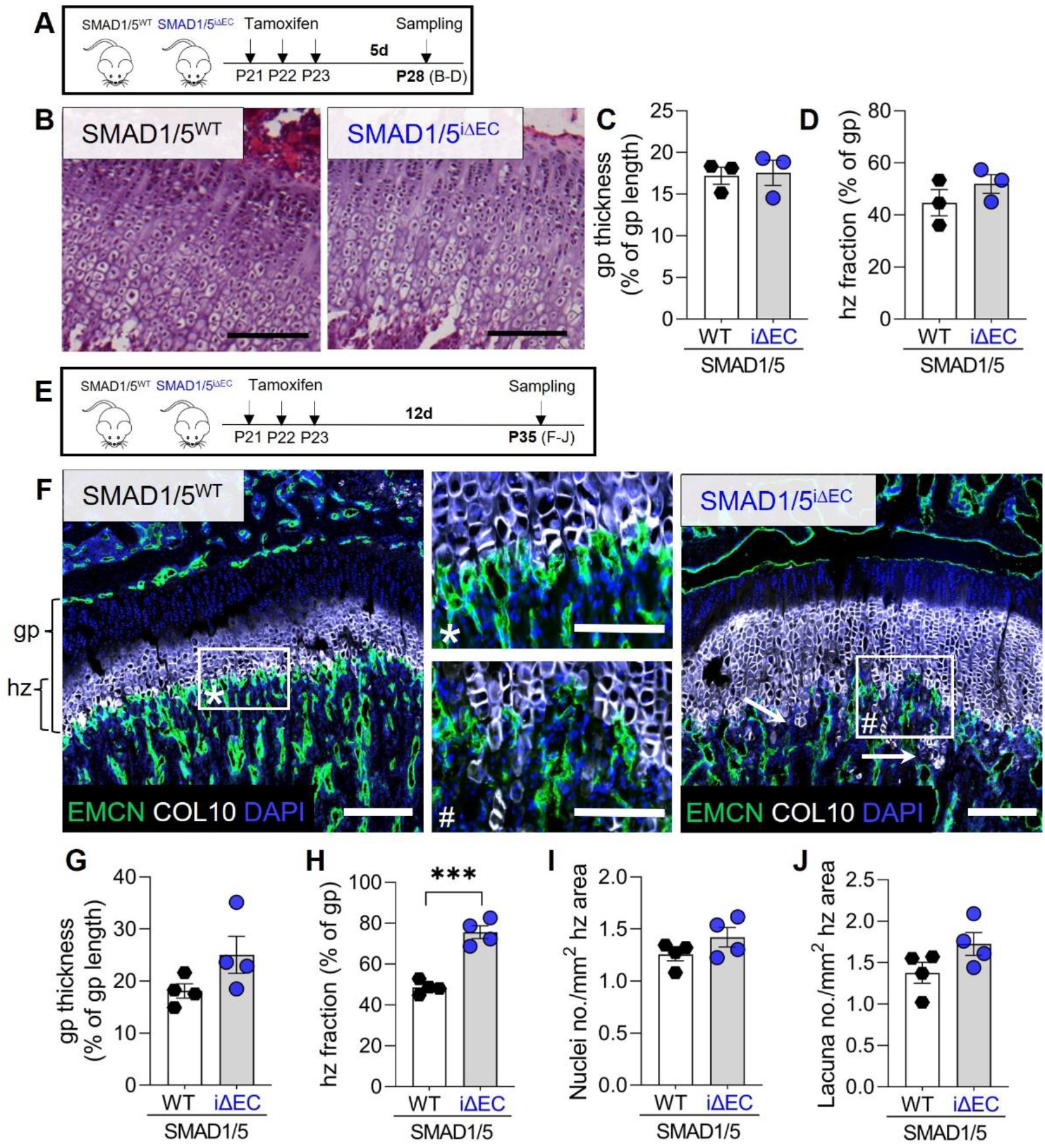
Integrity of the hypertrophic chondrocyte zone within the growth plate depends on functional adjacent type H vessel formation. (**A**) Tamoxifen treatment scheme. Mice were injected postnatal day 19-21 (P19-21) and samples were collected at P28. (**B**) Representative images of H&E staining at P28. Quantification of (**C**) growth plate thickness relative to the growth plate length and (**D**) hypertrophic zone fraction at P28 (n^WT^= 3; n^iΔEC^= 3). (**E**) Tamoxifen treatment scheme. Mice were injected postnatal day 19-21 (P19-21) and samples were collected at P35. (**F**) Representative images of EMCN, COL10 and DAPI staining in the epi-/metaphysis at P35. gp – growth plate; hz – hypertrophic zone; arrows indicate penetration of COL10 positive chondrocyte columns into the metaphyseal vascular area. Quantification of (**G**) growth plate thickness relative to the growth plate length, (**H**) hypertrophic zone fraction, (**I**) nuclei as well as (**J**) lacuna number in the hypertrophic zone area (n^WT^= 4; n^iΔEC^= 4). Bar graphs show mean ± SEM and individual data points. Two-sample t-test was used to determine the statistical significance; p-values are indicated with *\*\*\*p < 0.001*. All scale bars indicate 250 μm (**B, F**) or 125 μm (magnifications **F**).

### Endothelial SMAD1/5 signaling regulates endomucin expression and vascular maturation in diaphyseal sinusoidal (type L) capillaries

Type L vessels have sinusoidal structure, form by sprouting angiogenesis, and functionally couple with hematopoiesis in the bone marrow. To determine the role of postnatal SMAD1/5 signaling in type L vessel morphogenesis and maintenance, we performed histomorphometry for type L vessels at P28 (CD31^low^EMCN^low^; **Fig. 6A, B**) in the diaphysis (*4*). Endothelial SMAD1/5 depletion significantly increased the number and size of diaphyseal vascular loops (*p= 0.02;* **Fig. 6C**), confirming the CECT data. Endothelial-conditional SMAD1/5 depletion increased the EMCN^+^ area (mean difference= 22% ± 2%; *p< 0.001;* **Fig. 6D**) but differences in CD31^+^ area were not significant (mean difference= 5.8% ± 4.8%; *p= 0.27;* **Fig. 6E**), resulting in a significantly elevated EMCN/CD31 ratio (**Fig. 6F**). Consistently, *Emcn* mRNA was elevated 4-fold in the diaphyseal bone marrow area in SMAD1/5^iΔEC^ mice (*p< 0.001;* **Fig. 6G**) while differences in *Pecam* expression were not significant (*p= 0.12;* **Fig. 6H**). Since tip and stalk cell selection is guided by DLL4/Notch interaction with tip cells showing higher expression of DLL4, we analyzed mRNA expression of *Dll4* in the diaphysis and found significant higher expression in SMAD1/5^iΔEC^ mice (*p= 0.03;* **Fig. 6I**). We also observed significantly increased *Bmp6* mRNA expression in SMAD1/5^iΔEC^ mice (**Fig. 6J**). BMP6 has been shown to increase vascular permeability (*25*), therefore we next stained Ter119^+^ erythrocytes, revealing extensive extravascular blood cells, indicating vascular barrier dysfunction (**Fig. 6K**). Together, these data show that endothelial SMAD1/5 activity is essential to maintain the type L sinusoidal capillary phenotype in the diaphysis, with SMAD1/5 depletion inducing excessive formation of large vascular loops typically characteristic of type H vasculature, featuring elevated EMCN expression, ectopic tip cell formation, and vascular hyperpermeability.

**Figure 6.**
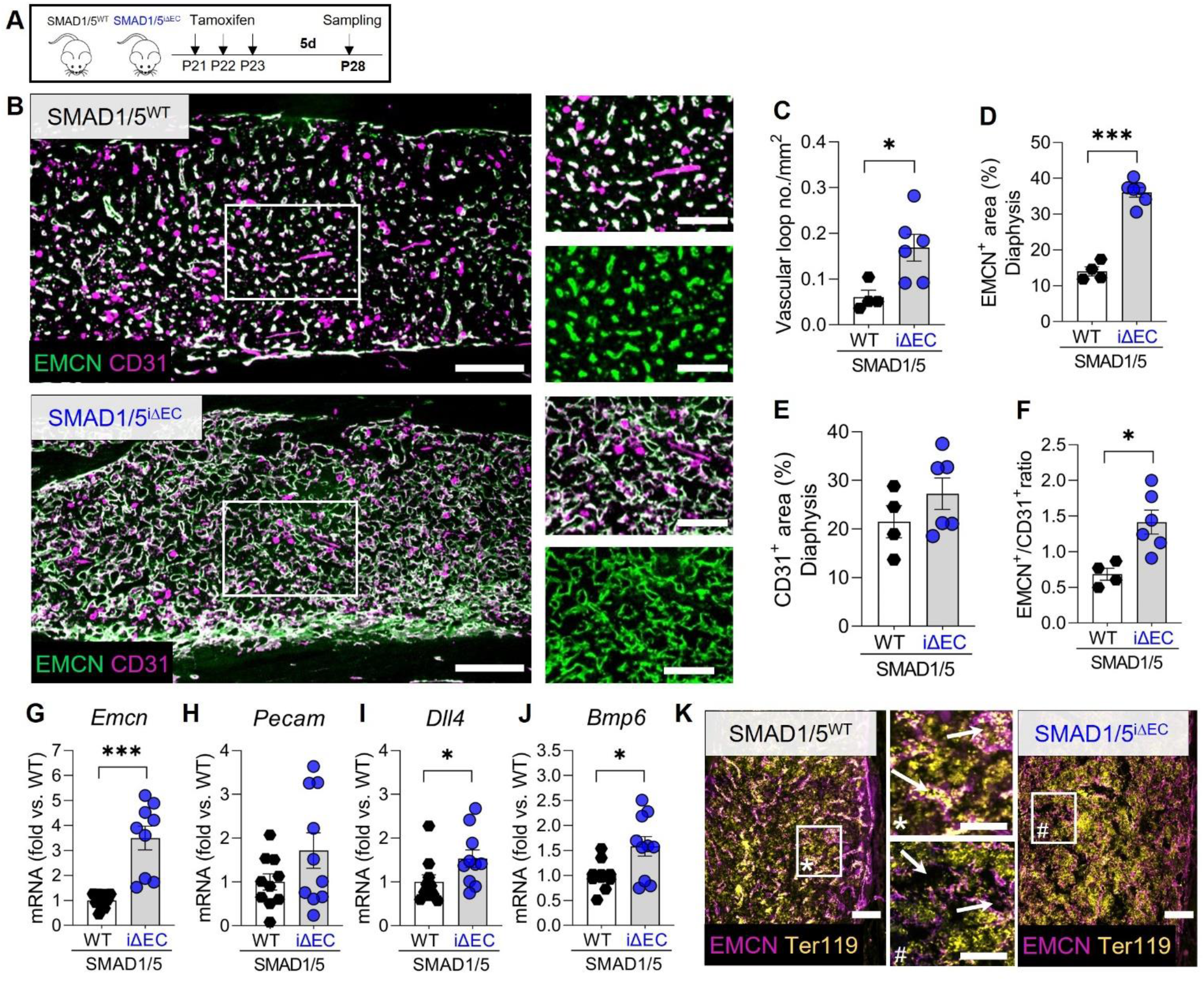
Endothelial SMAD1/5 promotes maturation and maintenance of diaphyseal sinusoidal (type L) capillaries. (**A**) Tamoxifen treatment scheme. Mice were injected postnatal day 19-21 (P19-21) and samples were collected at P28. (**B**) Representative images of EMCN and CD31 staining in the diaphysis (P28; n^WT^= 4; n^iΔEC^= 6). Quantification of (**C**) number of vascular loops per mm^2^, (**D**) relative EMCN^+^ and (**E**) CD31^+^ area and (**F**) EMCN^+^/CD31^+^ ratio (P28; n^WT^= 4; n^iΔEC^= 6). mRNA expression analysis of (**G**) *Emcn*, (**H**) *Pecam*, (**I**) *Dll4* and (**J**) *Bmp6* in the diaphysis (P28; n= 10). Bar graphs show mean ± SEM and individual data points. Two-sample t-test was used to determine the statistical significance; p-values are indicated with *\*p < 0.05; ***p < 0.001*. (**K**) Representative images of EMCN and Ter119 staining in the diaphysis (P28; n= 2) with magnifications highlighting intravascular* Ter119 staining or empty vessels# (additionally marked with arrows). All scale bars indicate 250 μm (**B**), 125 μm (**K**, magnifications **B**) or 62.5 μm (magnifications **K**).

### Endothelial SMAD1/5 activity is also required for metaphyseal and diaphyseal maintenance during early adolescence

Since the bone marrow vasculature undergoes continuous remodeling during postnatal and adolescent development, we next investigated the effects of EC-specific depletion of SMAD1/5 in more mature mice. Mice were injected with tamoxifen at P42 and samples were collected 7 or 14 days later, at P49 and P56 (i.e., 4 and 5 weeks post-weaning, respectively) (**Fig. 7B**, **K**). Analysis of type H vessels in the metaphysis at 7 days after endothelial-conditional SMAD1/5 depletion (P49) revealed impaired columnar structure, characterized by pronounced branching and network formation (**Fig. 7B**). Moreover, as in younger mice, 7 days of SMAD1/5 depletion reduced the number of anastomotic arches (*p = 0.02*) adjacent to the growth plate (**Fig. 7C**) and did not significantly alter EMCN^+^ area but significantly reduced CD31^+^ area (*p= 0.035;* **Fig. 7D, E**). Analysis of type L vessels in the diaphysis at 7 days after endothelial-conditional SMAD1/5 depletion (P49) revealed significantly increased diaphyseal vascular loop formation (*p= 0.047*) (**Fig. 7G**), as in younger mice and increased EMCN^+^ area (*p= 0.011*) with no differences in CD31^+^ area (*p= 0.21;* **Fig. 7H, I**). In line with this, the EMCN/CD31 ratio was elevated (*p= 0.021;* **Fig. 7J**). These alterations in type H and type L vasculature were qualitatively pronounced at P56 mice (14 days post-tamoxifen injection) (**Fig. 7L, M**). Moreover, OSX staining in the metaphysis indicated qualitatively reduced osteoprogenitor abundance at 14 days post-depletion, but this reduction was less dramatic than in young mice at P35 (**Fig. 7N,** cf. **Fig. 4I**). Similarly, endothelial SMAD1/5 depletion disrupted vascular loop formation at the chondro-osseous junction and disorganized cartilage septum, but without dramatic enlargement of the hypertrophic zone as observed before (**Fig. 7O,** cf. **Fig. 5F**). Together, these data support a model in which endothelial SMAD1/5 activity regulates type H vascular sprouting dynamics, maintains type L vascular stability, and coordinates growth plate remodeling and osteoprogenitor recruitment dynamics.

**Figure 7.**
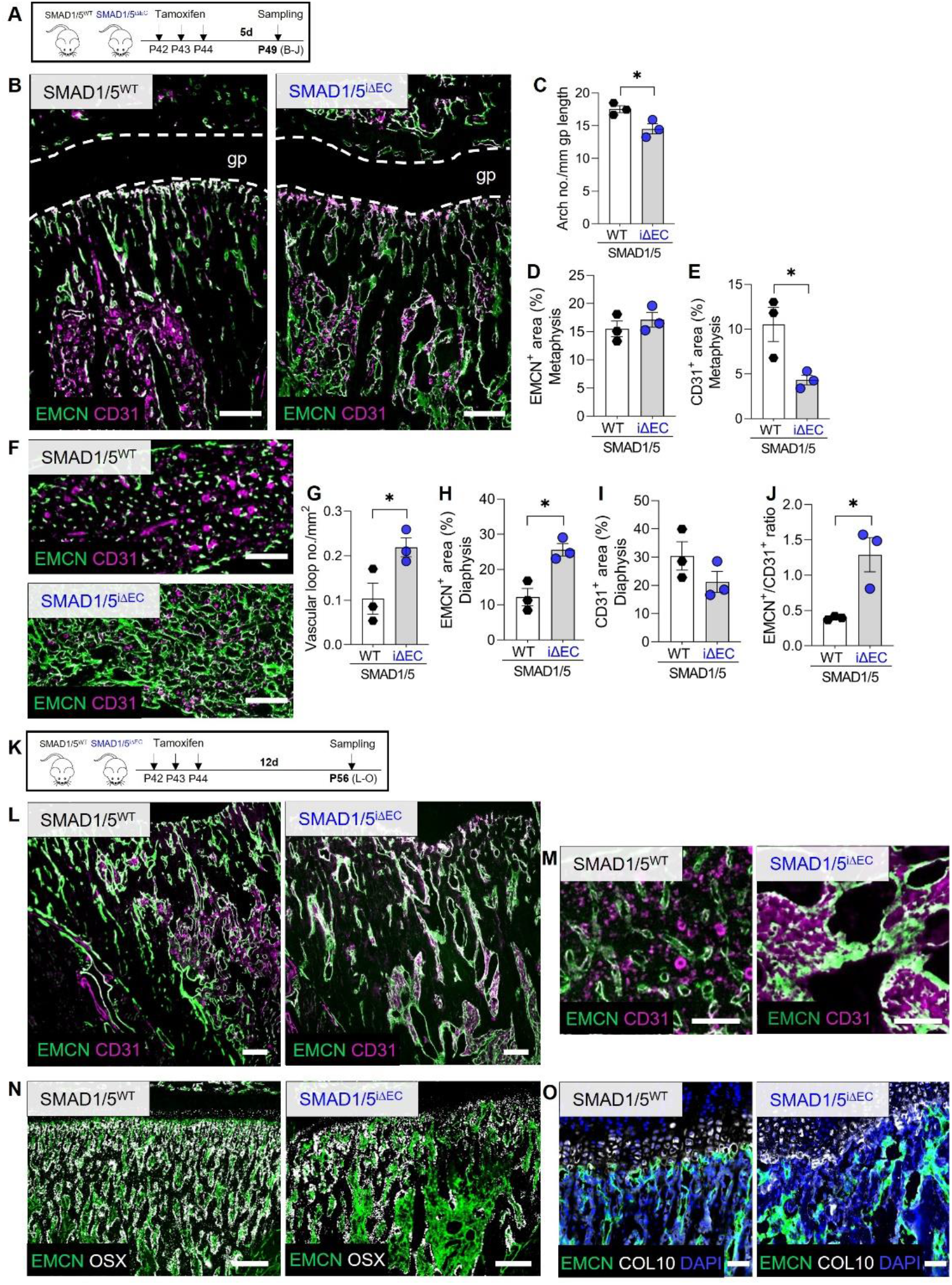
Endothelial SMAD1/5 maintains morphology and function of metaphyseal and diaphyseal capillaries during early adolescent. (**A**) Tamoxifen treatment scheme. Mice were injected postnatal day 42-44 (P42-44) and samples were collected at P49 (7 weeks - 4 weeks post-weaning). (**B**) Representative images of EMCN and CD31 staining in the metaphysis (P42; n^WT^= 3; n^iΔEC^= 3). (**B**) Quantification of (**C**) arch number, (**D**) relative EMCN^+^ and (**E**) CD31^+^ area. (**F**) Representative images of EMCN and CD31 staining in the diaphysis (P49; n^WT^= 3; n^iΔEC^= 3). Quantification of (**G**) number of vascular loops per mm^2^, (**H**) relative EMCN^+^ and (**I**) CD31^+^ area as well as (**J**) EMCN^+^/CD31^+^ ratio. Bar graphs show mean ± SEM and individual data points. Two-sample t-test was used to determine the statistical significance; p-values are indicated with *\*p < 0.05*. (**K**) Tamoxifen treatment scheme. Mice were injected postnatal day 42-44 (P42-44) and samples were collected at P56 (8 weeks - 5 weeks post-weaning). Representative images of (**L**) EMCN and CD31 staining in the metaphysis or (**M**) diaphysis (P56; n^WT^= 3; n^iΔEC^= 2). Representative images of (**N**) EMCN and OSX or (**O**) EMCN, Col X and DAPI staining in the metaphysis (P56; n^WT^= 3; n^iΔEC^= 2). All scale bars indicate 250 μm (**B**, **F**, **L**, **N**), 125 μm (**M**) or 62.5 μm (**O**).

## Discussion

Here, we show that endothelial SMAD1/5 activity sustains skeletal vascular morphogenesis and function and coordinates growth plate remodeling and osteoprogenitor recruitment dynamics during juvenile and adolescent bone growth (**Fig. 8**). We found that endothelial cell-conditional SMAD1/5 depletion in juvenile mice caused hypervascularity in both metaphyseal and diaphyseal vascular compartments, resulting in altered cancellous and cortical bone formation. Short- and long-term SMAD1/5 depletion, in both juvenile and adolescent mice, induced excessive sprouting, disrupting the columnar structure of the type H metaphyseal vessels and impaired anastomotic loop morphogenesis at the chondro-osseous junction. SMAD1/5 depletion progressively arrested osteoprogenitor recruitment to the primary spongiosa and, in the long term, impaired growth plate resorption. Finally, in the diaphyseal sinusoids, endothelial SMAD1/5 activity was necessary to maintain the type L phenotype, with SMAD1/5 depletion inducing excessive formation of large vascular loops typically characteristic of type H vasculature, featuring elevated endomucin expression, ectopic tip cell formation, and vascular hyperpermeability. Together, these data show that SMAD1/5 signaling in the endothelium preserves skeletal vessel structure and function and couples angiogenesis to osteogenesis during both juvenile and adolescent bone growth.

**Figure 8.**
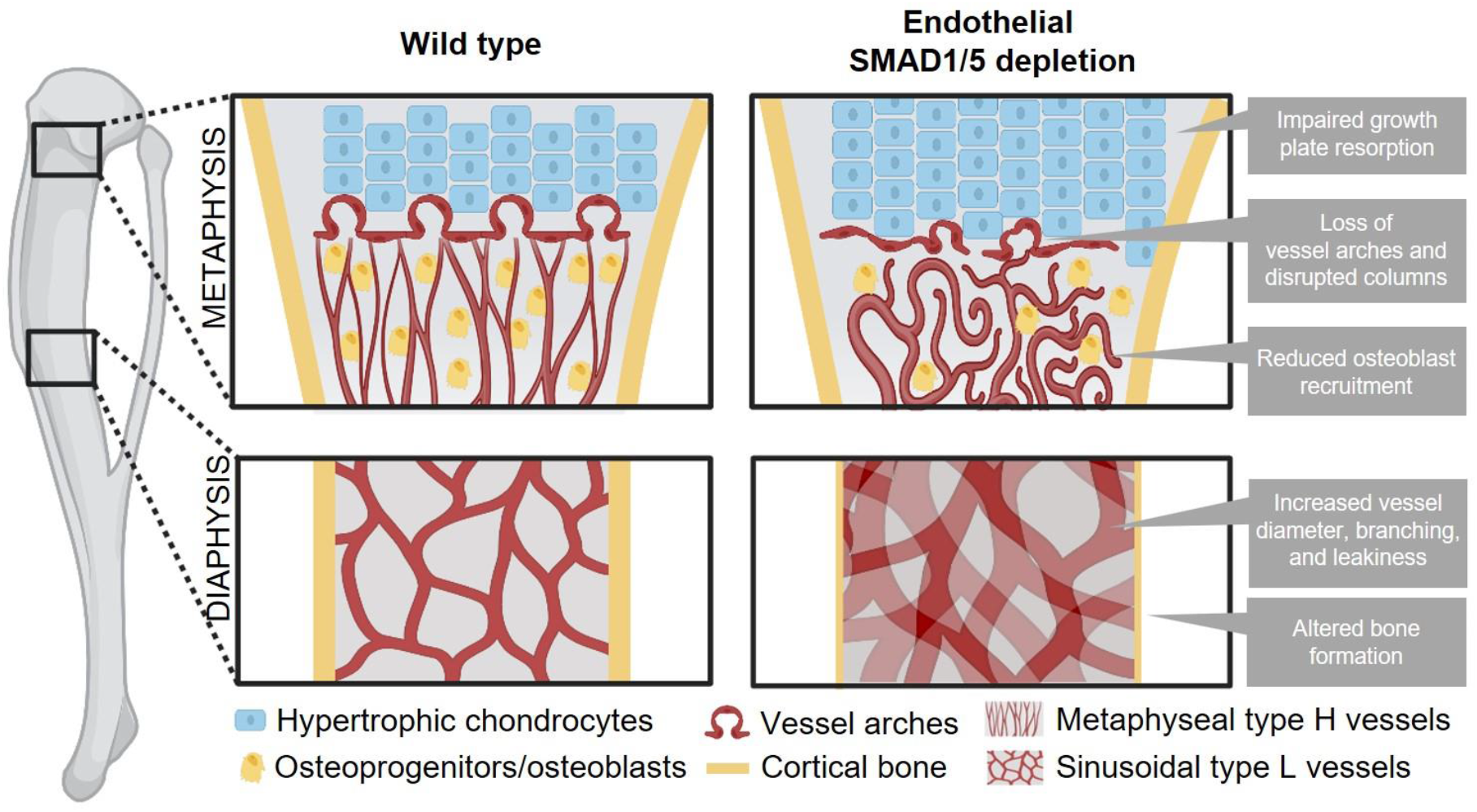
Graphical summary on effects of endothelial SMAD1/5 depletion on metaphyseal and diaphyseal vessel and bone formation during juvenile and early adolescent long bone growth. Illustration has been created with BioRender.com.

Juvenile skeletal growth requires dynamic adaptation of bone formation accompanied by a substantial adjustment of the bone vasculature. Distinct vascular morphology has been described starting at postnatal day 6 with metaphyseal capillaries (type H vessel) showing a column-like structure and diaphyseal capillaries (type L vessel) forming a sinusoidal network (*26*). Murine long bone growth evolves rapidly until P14, reaching a steady growth phase between P14 and P42 (*27, 28*). This is in accordance with the already described rapid decline of the Type H vessel over the first 4 weeks postnatally (*4*). We found that EC-specific SMAD1/5 depletion at P21 resulted in a substantial enlargement of the diaphyseal vasculature with less dramatic changes in the metaphyseal vasculature at P28. This observation suggests that endothelial SMAD1/5 signaling i) directs morphogenesis of both metaphyseal and diaphyseal vessels during juvenile long bone growth and ii) maintains vascular stability, contributing to the transformation of type H to type L vessels in the diaphysis from between P14 and P28. These changes to the vasculature altered bone formation. Particularly corticalization was impaired after short term depletion of endothelial SMAD1/5 activity (P28). Long bone growth features endochondral bone formation at the growth plate, leading to trabecular bone formation, which coalesces at the metaphyseal cortex with bone lining-cell intramembranous ossification, initiating the corticalization process (*29*). Depending on age and the location, the cortical shell is formed from corticalizing trabeculae in a vigorous process (*30*). Thus, the significant decrease cortical area and porosity upon EC-specific SMAD1/5 depletion indicates disturbed corticalization which is especially driven by Osterix-expressing osteoblasts (*31*) and therefore, indicates alterations in angiogenic-osteogenic coupling.

At the chondro-osseous junction, Osterix-expressing osteoprogenitors spatially localize with type H endothelium and mediate angiogenic-osteogenic coupling by multiple pathways, including Notch signaling (*4, 32*). Comparable to the shape maintaining function of endothelial Notch signaling (*32*), we show here that SMAD1/5 activity in type H endothelium is crucial to maintain their archetypal columnar structure and new arch formation in both juvenile (P21) and adolescent (P42) bones. DLL4-Notch signaling is responsible for tip and stalk cells competence in the metaphysis and is driven by crosstalk between ECs and chondrocytes (via VEGF, Noggin) (*32*). We demonstrated previously that during mouse embryonic development, Notch and SMAD1/5 signaling synergize to balance selection of tip and stalk cells in vascular sprouting (*19*). Synthesizing these findings with our present results, supported in part by general bulk elevation of Notch-related gene expression (*Dll4*) in SMAD1/5 SMAD1/5^iΔEC^ mice, we hypothesize that the alterations in type H vessel angiogenesis in the metaphysis result from the disrupted Notch/SMAD1/5 synergy in the bulging vessels. In addition, BMP2/6/7, which signal through SMAD1/5, are abundant in bone (*10, 12*). These ligands guide endothelial tip cell competence via type I receptors (ALK2, ALK3, ALK6), in conjunction with BMP type II receptor (*33*), suggesting that bulging angiogenesis by type H vessels in the metaphysis may be regulated by BMP-SMAD signaling. Consistently, we observed profound disruption of angiogenic-osteogenic coupling in the metaphysis, with reduced *Sp7* mRNA expression at 7 days post-depletion and near complete abrogation of OSX-expressing cells in the metaphysis after 14 days. Since OSX^+^ cells substantially expand during the first 4 weeks postnatally in the metaphysis (*26*), these time-dependent findings indicate the requirement of continued endothelial SMAD1/5 activity in osteoprogenitor survival and recruitment during endochondral bone growth. Further studies are required to investigate the angiogenic-osteogenic crosstalk mechanisms and the fate of the osteoprogenitors upon endothelial SMAD1/5 depletion.

Endochondral bone formation at the chondro-osseous junction requires neovascular invasion and growth plate remodeling. Previous studies reporting enlargement of the growth plate, especially the hypertrophic zone, upon disruption of the growth plate-adjacent vasculature by inhibition of VEGF signaling (*34, 35*) or endothelial MMP9 depletion (*24*). Consistent with these data, we found that EC-specific SMAD1/5 depletion resulted in a significant enlargement of the hypertrophic zone of the growth plate. Therefore, the dysmorphogenesis of the anastomotic type H endothelium arches at the chondro-osseous junction upon EC-specific SMAD1/5 depletion could be the reason for the enlargement of the hypertrophic chondrocyte zone. This is further supported by the observation in retinal angiogenesis that BMP4-SMAD1/5 signaling regulates endothelial MMP9 function (*36*). Based on our finding that the number of DAPI^+^ cells and chondrocyte lacuna in the COL10^+^ area were only slightly increased, we propose that EC-specific SMAD1/5 inactivity affected the removal of cartilage matrix and the transition from hypertrophic chondrocytes to bone rather than chondrocyte hypertrophy, *per se* (*37*).

Growth plate remodeling and endochondral ossification are mediated by type H vessels, but the long bone diaphysis is populated by sinusoidal type L vessels. These vessels have been proposed to be maintained in a homeostatic and quiescent state with relatively slower physiological remodeling (*4*). Type L vessels likely emerge through maturation of type H capillaries (*4, 26*). While there is evidence suggesting constant remodeling and volume adaptations in the diaphyseal capillary network (*38*), the dynamics and underlying mechanisms are mostly unknown. We previously showed that EC-specific depletion of SMAD1/5 during early postnatal retinal angiogenesis resulted in arteriovenous malformations, a reduced number of tip cells, and hyperdensity in the retinal vascular plexus (*20*). These findings mirror the diaphyseal vessel changes, characterized by significant hyper-density and aberrant vascular loop formation. We observed progressive emergence of diaphyseal vessels with characteristics of type H vessels (i.e., significantly elevated EMCN and CD31 expression) upon EC-specific depletion of SMAD1/5. Tip and stalk cell selection during sprouting angiogenesis is guided by DLL4/Notch interaction, with tip cells showing higher expression of DLL4 (*32*). Previously, we found that endothelial SMAD1/5 specifically regulates Notch-mediated tip cell formation in the E9.5 mouse hindbrain (*19*), consistent with our observation here of significantly elevated mRNA expression of *Dll4* and *Bmp6* in the diaphyseal bone marrow. Thus, the hyper-dilatation of the diaphyseal vasculature may be a result of pronounced bulging angiogenesis (sprouting) in the type L vessels and progressive conversion to the type H phenotype, including an increase in tip-like endothelial cells, upon cessation of SMAD1/5 signaling. Vascular homeostasis, quiescence, and maturation are controlled by BMP9/10 signaling via ALK1-BMPR2 complexes activating SMAD1/5 (*39*). BMP9/10-ALK1-SMAD1/5 signaling may therefore modulate homeostatic signaling in type L vessel maturation and phenotypic maintenance. Together, these findings suggest a central role of endothelial SMAD1/5 in maintenance of sinusoidal vascular homeostasis.

A comparable phenotype of hyper-dilated and functionally leaky vessels has been described in mouse embryos with a global loss of the BMP receptor Activin receptor-like kinase 1 (ALK1) or adult mice with an endothelial-specific ALK1 knockout (*40, 41*). Genetic defects in ALK1 signaling cause the autosomal dominant vascular disorder, hereditary hemorrhagic telangiectasia (HHT), which causes arteriovenous malformations (AVM) and vessel wall fragility, resulting in a risk for fatal hemorrhage in human patients (*42*). Arteriovenous malformations in the bone marrow have been described (*43*).

In conclusion, this study identifies SMAD1/5 signaling in endothelial cells as an essential regulator of vascular formation, maturation, and homeostasis in long bones, and as a mediator of angiogenic-osteogenic coupling. Our findings underline the importance of functional BMP-SMAD signaling in long bone vasculature during bone growth and may inform clinical management of congenital diseases like HHT (*43*) and the development of new therapies for enhancing vascularized bone repair and regeneration (*13, 44–46*).

## Limitations

This study had limitations: Based on our experimental design, we cannot draw conclusions on the functional role of SMAD1/5 in angiogenic-osteogenic coupling during embryogenesis and early postnatal bone development. Further studies will be necessary to dissect these early timepoints which exhibit more rapid cellular dynamics and unique cell populations compared to juvenile and adolescent skeletal formation. We have previously reported that a constitutive EC-specific depletion of SMAD1/5 activity is embryonically lethal (*19, 20*), so continued study using the inducible system is warranted. In our study, we found that the serious malformations in the vascular system precluded analysis of samples collected at later timepoints after tamoxifen induction (14 days). This resulted in lower sample sizes in the analysis of P35 or P56. In addition, our experimental approach designed to detect differences according to sex as an independent variable, but both sexes were included in the study and equal distribution of data did not provide evidence of sexual dimorphism.

## Materials and Methods

### Breeding strategy and housing

Mice were housed and bred in the Animal Facility at KU-Leuven (Belgium) and all animal procedures were approved by the Ethical Committee (P039/2017, M007, M008). Breeding was performed as described previously (*20*). In detail, homozygous mice caring the Smad1/Smad5 floxed alleles (Smad1^fl/fl^;Smad5^fl/fl^) were paired with endothelium-specific tamoxifen-inducible Cre mice expressing (Cdh5-CreERT2^tg/0^). Subsequently, dams (Smad1^fl/fl^;Smad5^fl/fl^) were crossed with the obtained Cdh5-CreERT2^tg/0^;Smad1^fl/+^;Smad5^fl/+^ mice. The resulting Cdh5-CreERT2^tg/0^;Smad1^fl/fl^;Smad5^fl/fl^ pups were injected intraperitoneally with tamoxifen (500 μg; Sigma Aldrich) at i) postnatal day 19, 20 and 21 (P21) or ii) postnatal day 42 (6-week old) to create EC-specific double knockout pups(SMAD1/5^iΔEC^). Pups were killed at P28 or P35, or at P49 (7-week old) or P56 (8-week old). Mice have a mixed background of CD1 and C57Bl6. All experiments were conducted using Cre-negative littermate controls. Genotyping of recombined alleles was done after sample collection as previously described (*19*). For breeding, mice were housed in pairs (one male and one female) in IVC Eurostandard Type II clear-transparent plastic cages (two animals per cage) covered with a wire lid and built-in u-shaped feed hopper and closed with a filter top in a SFP barrier facility. Weaning was performed at an age of approx. 3 weeks while littermates were housed together with 5 mice in Eurostandard Type II cages and transferred to a semi-barrier facility with IVC cages. As bedding material, fine wood chips and Nestlets for nesting were provided as well as plastic houses for environmental enrichment. The room temperature was constant in both facilities between 20 and 22°C and a 12/12-h light/dark cycle with lights on at 0700 hours and off at 1900 hours. Mice received standard diet and tap water *ad libitum*. Mice were killed by cervical dislocation. Male and female mice were used for investigations and sex-specific differences were not analyzed. All experiments and analyses were conducted with samples from at least 3 different litters/experiments.

### Contrast-enhanced microfocus X-ray computed tomography

Right tibias from SMAD1/5^WT^ and SMAD1/5^iΔEC^ mice (n= 10 per group) were collected and fixed in 4% paraformaldehyde (PFA; Sigma Aldrich) in PBS overnight (12h) at 4°C. Samples were stored in PBS at 4°C until further use to allow for consistent staining of all samples with an X-ray contrast-enhancing staining agent (CESA). The distal part of the tibia was cut to open the shaft and allow for uniform distribution of the CESA solution. Samples were stained for 1 week with a Hafnium-substituted Wells-Dawson polyoxometalate (POM) solution (35 mg/ml PBS) at 4°C under constant shaking as established previously (*47*). High-resolution microfocus computed tomography (μCT) imaging was performed with a GE NanoTom M (GE Measurement & Control) at 60 kV and 140 μA, with a 0.2 mm filter of aluminum and a voxel size of 2 μm. The exposure time was 500 ms, and 2400 images were acquired over 360° using the fast scan mode, resulting in 20 minutes acquisition time. During reconstruction (Datos-x, GE Measurement & Control), a beam hardening correction of 7 and a Gaussian filter of 6 was used. Detailed structural analysis of all datasets was performed using CTAn (version 1.16) and DataViewer (both Bruker Corporation). Volumes of interest (VOIs) of 301 images (0.6 mm) were analyzed in the metaphysis and diaphysis, respectively. To determine the starting point of the metaphyseal and diaphyseal area, the image displaying the middle part of the growth plate was manually determined (GP). The start of the metaphyseal VOI was determined 300 images downstream of the GP, while the diaphysis started 100 images under the end of the metaphyseal area, representing the transition zone between meta- and diaphysis. Thresholding for binarization of the vessels was manually performed based on the histogram, while for bone binarization, automatic Otsu thresholding was applied. Manually drawn ROIs of the bone area (outer cortical surface) were specified with the ROI shrink-wrap tools stretching over holes with a diameter of 60 pixels and independent objects were removed using the despeckle tool. For analysis, the provided task set for 3D analysis was employed including analysis of structure separation distribution for the vessels. 1,000 images of exemplary samples were used for 3D rendering and visualization (CTvox; version 3.2.0; Bruker Corporation).

### Classical 2D histology and immunofluorescence

For immunofluorescence, tibias were fixed in 4% PFA in PBS for 6-8h at 4°C. Samples were cryo-embedded (SCEM medium, Sectionlab) after treatment with an ascending sucrose solution (10, 20, 30%) for 24h each. Sectioning was performed using a cryotape (Sectionlab) and sections were stored at −80°C until further use. For immunofluorescence staining, sections were airdried for 10 min before being hydrated in PBS (5 min). Blocking solution contained 10% donkey or goat serum in PBS (30 min) and antibodies were diluted in PBS/0.1% Tween20/5% donkey or goat serum (Sigma Aldrich). The following primary antibodies and secondary antibodies were used (staining durations are individually provided): pSMAD1/5 (Cell signaling; clone: D5B10; 1:100; incubation over night at 4°C); CD31/PECAM-1 (R&D Systems; catalog number: AF3628; 1:100; 2h at RT - room temperature), EMCN (Santa Cruz; clone V.5C7; 1:100; 2h at RT), COL10 (Abcam; catalog number: ab58632; 1:100; 2h at RT); OSX (Abcam; catalog number: ab209484; 1:100; 2h at RT), Terr119-APC (Biolegend; 116223); all secondary antibodies were purchased from Thermo Fisher Scientific and used at an 1:500 dilution for 2h at RT if not stated otherwise: goat anti-rat A647 (A-21247), donkey anti-goat A568 (A-11057), goat anti-rat A488 (A-11006), goat anti-rabbit A647 (A-27040), goat anti-rabbit A488 (Abcam; ab150077; 1:1,000). DAPI (1:1,000; Sigma Aldrich) was added during the last washing step and sections were covered with Fluoromount_GT (Thermo Fisher Scientific), and a cover slip. Images were taken with a Keyence BZ9000 microscope (Keyence), a Zeiss LSM880 or an AxioScan (both Carl Zeiss Microscopy Deutschland GmbH) and image quantification was performed in a blinded manner using the Fiji/ImageJ software. The area of interest was manually created and managed with the built-in ROI-Manager. Arches, nuclei, and lacuna were counted manually, and areas (%) were determined with the thresholding tool.

### RNA analysis

For RNA analysis, left tibias from mice used for CECT analysis were treated with RNAlater (Qiagen) and stored at −80°C until further use. Separation of the metaphysis and diaphysis was done by cutting underneath the growth plate. Bulk samples including bone and bone marrow were cryo-pulverized and resuspended in 1 ml ice-cold TriFast (VWR International) and carefully vortexed (30 sec). A volume of 200 μl 1-bromo-3-chloropropane (Sigma Aldrich) was added and the mixture was incubated for 10 min at room temperature before centrifugation (10 min at 10.000 *x g*). The top aqueous phase was collected for RNA isolation using the RNeasy Mini Kit (Qiagen) following the manufacturer’s instructions. Purity of the RNA was analyzed via Nanodrop; RNA integrity and quality were verified via Fragment Analyzer. cDNA synthesis was performed using the TaqMan Reverse Transcription Reagents (Applied Biosystems; 0.5 μg/μl RNA concentration) and DyNAmo Flash SYBR Green qPCR Kit (Thermo Fisher) was performed at a Stratagene Mx3000P (Agilent Technologies) with the following protocol: 7 min initial denaturation at 95 °C, 45 to 60 cycles of 10 s denaturation at 95 °C, 7 s annealing at 60 °C and 9 s elongation at 72 °C (duplicates per gene). CT values were normalized to *Hprt*(Housekeeper); as second control *18s rRNA* was carried along. Primer were design using NCBI and Blast, tested and verified via Gel electrophoresis.

### Statistical analysis

GraphPad Prism V.8 was used for statistical analysis. Data was tested for Gaussian distribution according to D’Agostino-Pearson omnibus normality test and homoscedasticity. When parametric test assumptions were met the Student’s t-test was used to compare two groups. In case of failing normality testing, data were log-transformed, and residuals were evaluated prior to parametric testing on log-transformed data. A *p value* <0.05 was considered statistically significant. Sample sizes are indicated in the figure legends. Data are displayed with error bars showing mean ± SEM and individual samples in a bar graph.

**Table 1:**
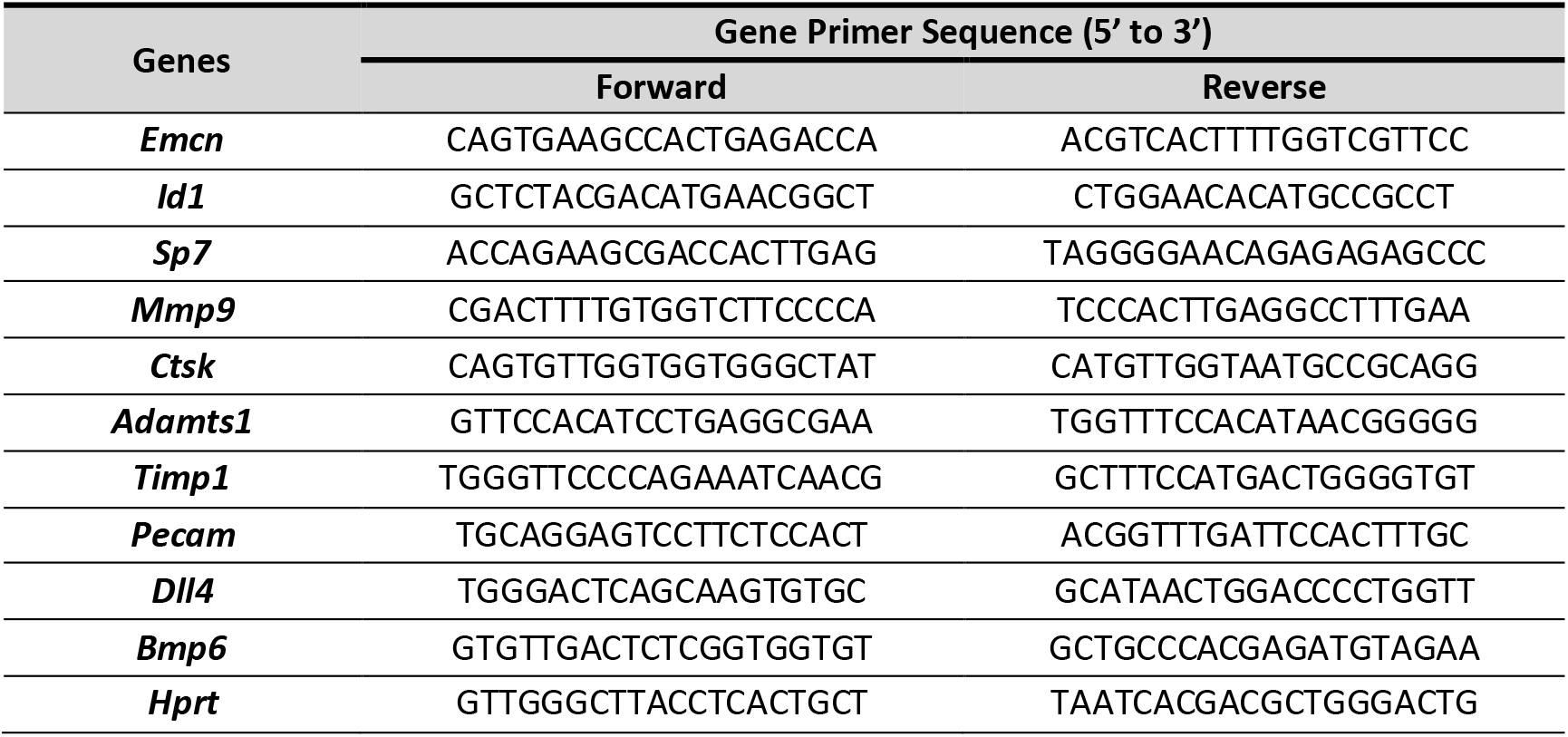
Primer sequences.

## H2: Supplementary Materials

## General

The authors would like to thank all members of the Zwijsen and Boerckel labs for constructive discussions on the research presented in this study. Furthermore, we like to thank Carla Geerome for assistance with the CECT scan.

## Author contributions

A. Lang, A. Benn, A. Zwijsen, and J.D. Boerckel conceived and supervised the research. A. Benn designed and performed animal experiments. A. Lang, A. Wolter and J. Collins performed analysis. G. Kerckhofs and T. Balcaen supervised and conceived CECT scans and generated the CESA. A. Lang, A. Benn, A. Zwijsen and J. D. Boerckel wrote the paper. All authors revised the manuscript.

## Data and materials availability

All data associated with this study are present in the paper or the Supplementary Materials.

## Supplementary Figures

**Figure S1.**
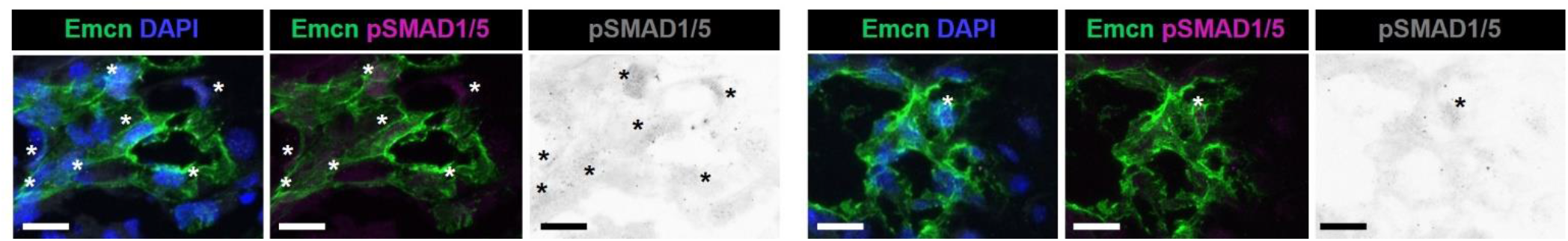
Reduction of endothelial phosphoSMAD1/5-positive ECs in the bone marrow of SMAD1/5^iΔEC^ mice. Mice were injected postnatal day 19-21 (P19-21) and samples were collected at P28. Figures show representative images of phospho(p)SMAD1/5 staining in EMCN positive endothelial cells in the diaphysis (P28; representative for n^WT^= 4; n^iΔEC^= 6). Scale bars indicate 20 μm. Asterisks indicate positive staining.

**Figure S2.**
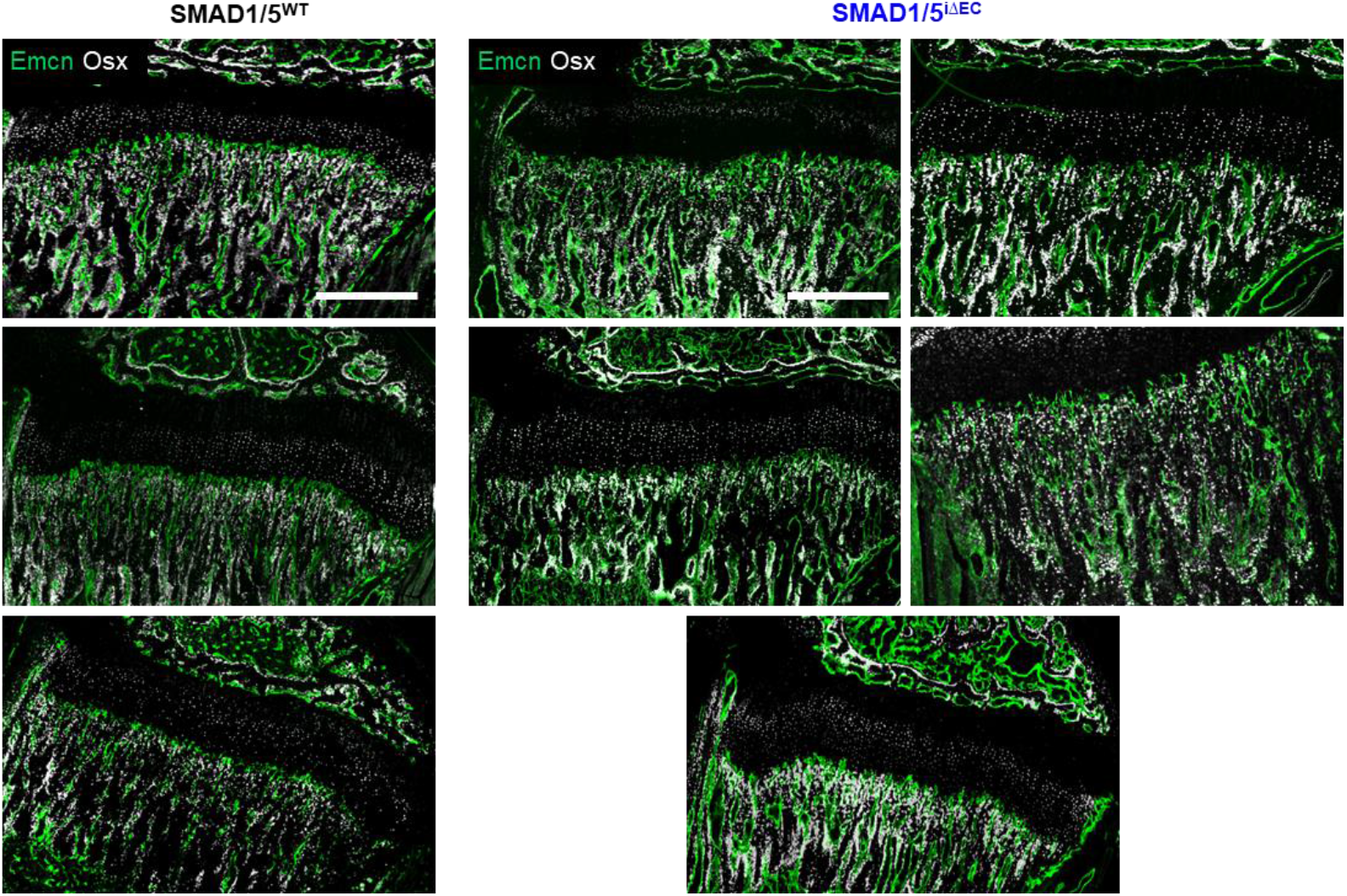
Additional images on co-localization of active osteoprogenitors in the metaphyseal area. All images of EMCN and OSX staining are in the tibial metaphysis (P28; n^WT^= 3; n^iΔEC^= 5). Scale bars indicate 500 μm.

**Figure S3.**
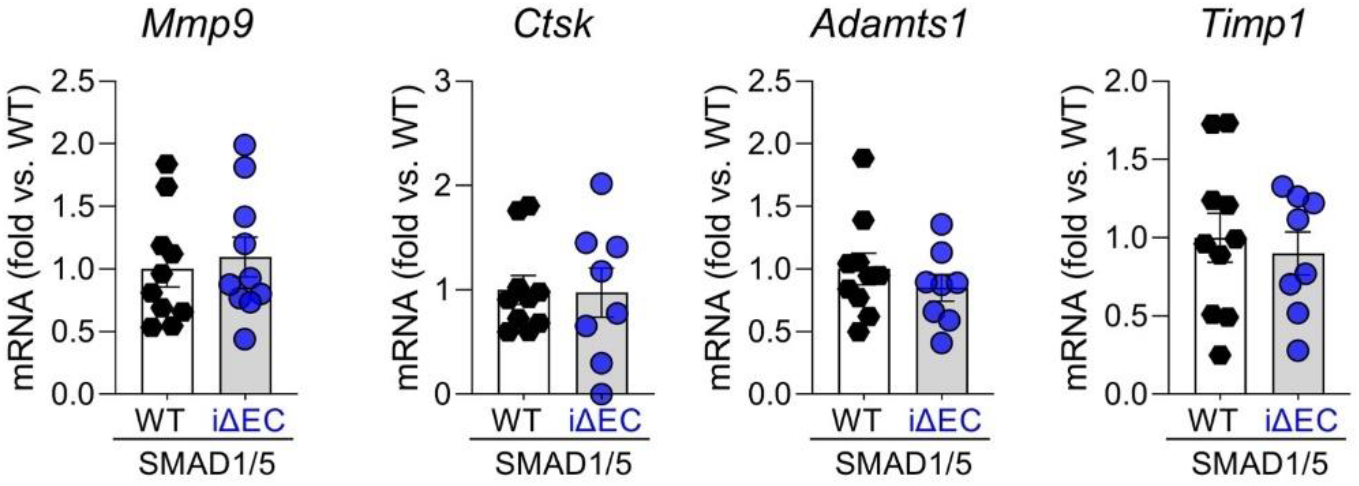
mRNA expression analysis of *Mmp9, Ctsk, Adamts1* and *Timp1* in the epi-/metaphysis. (P28; n= 10).

